## Supplemental Material for "Fine-tuning FAM161A gene augmentation therapy to restore retinal function"

### Supplemental Figure 1

**A**

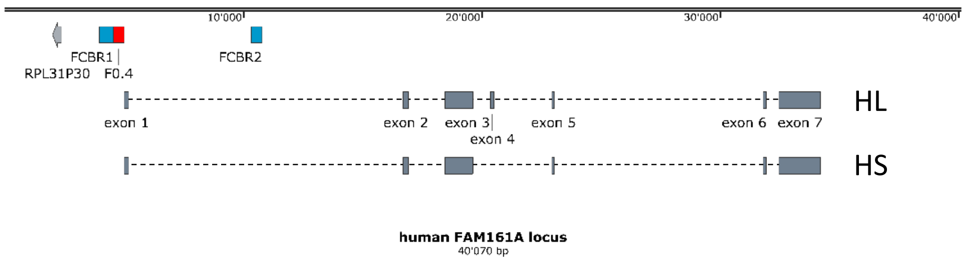


**B**

FCBR1 (enhancer, 607 bp)

GGCTCAGACCTTAGAGACGGGTTCTCGACGATGTGCAGCGCATAACGAGGAAAATACTGACCGGAGGGGGCCGGTCGTAGGGCTGGGG
AGCAGGAGGAGGGCAAAGGATAGAGAGGGGGTAGATAGGGAGTAGATAGGGGGCGGCAGCCTCGGCCTCAGTAAGCAGGGGCTCAGG
GAGCAGATGTCTGAGACTGCACCACGTCCGCTATTTTGAGGTTTGGCTTCTCTTACTGCCACGTGGTGCAGAGGGAAGGCGCCATTTGAT
GCGTCCAAATCCCGAGTGGGAAGAGAAGATACGGATGAGGGAGATGGATGTGTCTGTGGTGACTCTTGGGCACTTGTGCCGTTCAATCCT
GTCACTCTCAAGCTGAAGAACCCAAAGCAAACGCCAGGTTTACAATACCTGCACTATCACTC**ATTA**GTGGGAGGGTCAGGCGCCACCCAA
CTCTGGGCTTTTATTGGTCATGTCAAGAAAAAAGCGAGAGGGACAACTGGGAGCCTCCTTCTGATTGGTGGATTCCTGCTAGCCCTCAAGT
CAATCAGAGTAGAATGTACCGCCCTCCTTCCCACTCTGGCCCCCGCGGGAGACATGTTAGGTAGCGGGACG

FCBR2 (enhancer, 457 bp)

CTCTCTCATTCTTTACAAATTCAGCGAAAAAAATTTTTTCAAAA**ATTA**CAGGTTGCAGATGAAACTATATAGGCTGCAGCACAAA**TAAT**TCTT
CATAGGTGCTG**ATTA**C**TAAT**AAAGCAACAGGTGCTGGAAGGATTC**ATTA**TCTAAATGGGTCAAGCCAATCTAGGGAAG**ATTA**TCCTC**TAAT**
CTGGTGGCTGCATTTACAACATTCCACAAAAAGCCG**ATTA**AAGGGAGAGGATGCTATTTTAGGCTCTTGGAAAAA**TAAT**GTCTATGGCCTA
GGTTATTTTGTATACTAAATATTTTCTCAGTGGAGACA**ATTA**AAG**ATTA**A**ATTA**GAATTGGAGGTCATTTACTAGCCATTCATCTGTTTTAGGT
TAAACACTGTGAGATCAGTTTATCTGAAAAGTTTCGGCTAGCGTTAACAGT**ATTA**AGCTGAAAACAGAATGGATCCAGTGCAAAATCA

F0.4 (promoter, 460 bp)

CCAACCTTGTCACTCTTCTGAGACCGCCACCCAGTAACACGCCCTGCTTTCCTCCGCCGCCGCCGTCTCCTCCATCAACCGTCTCCTTCT
AACATGTTATTGCAGT**TAAT**AAAAAAGCGTTCTTTTCTTCACAAAAGCAGTTGTTTCATTGTG**TAAT**GTGAGAAGTCTGCGTTCAGGCGGTTA
GACGCAAAGGAGTGGAATAAGTGCCCTGTTCAGCCTGCCAACCTTCGGAACCAGCCAACAGACTCGGTACTGACTGCCATGATCCAGTAA
GCACCCGACTCGGGTCTCTGGCAGCCTGAGTGGATTTTGCGTGACTTTTGGTCCCTGGCGGCTCCTGTAGCGTCCCCAGTTACGGCGCC
CATAGCAACCGGCTCCCTAGCTAGGCGCCCCCGGGTTGCCAGGGGCCGCACCAGCTTTCCCGCCCCGGGCCAGCGCAGGCGCTCAGG
CCTCGGAGGCG

**Supplemental Figure 1. FAM161A-derived regulatory regions.** (**A**) Genomic region of human FAM161A is presented with the localisation of FCBR1, FCBR2 and F0.4 elements used to drive expression of HL or HS isoform via AAV vectors. The map corresponds to 1 to 40000 bp of genbank sequence NG028125.2 comprising the loci RPL31P30 and FAM161A. Gray box correspond to exon sequence, red box to the proximal FAM161A promoter and blue box to FCBR1 and FCBR2 region. (**B**) Sequences of FCBR1, FCBR2 and F0.4 elements with potential crx binding sites in bold and grey boxes.

**Supplemental Figure 2**

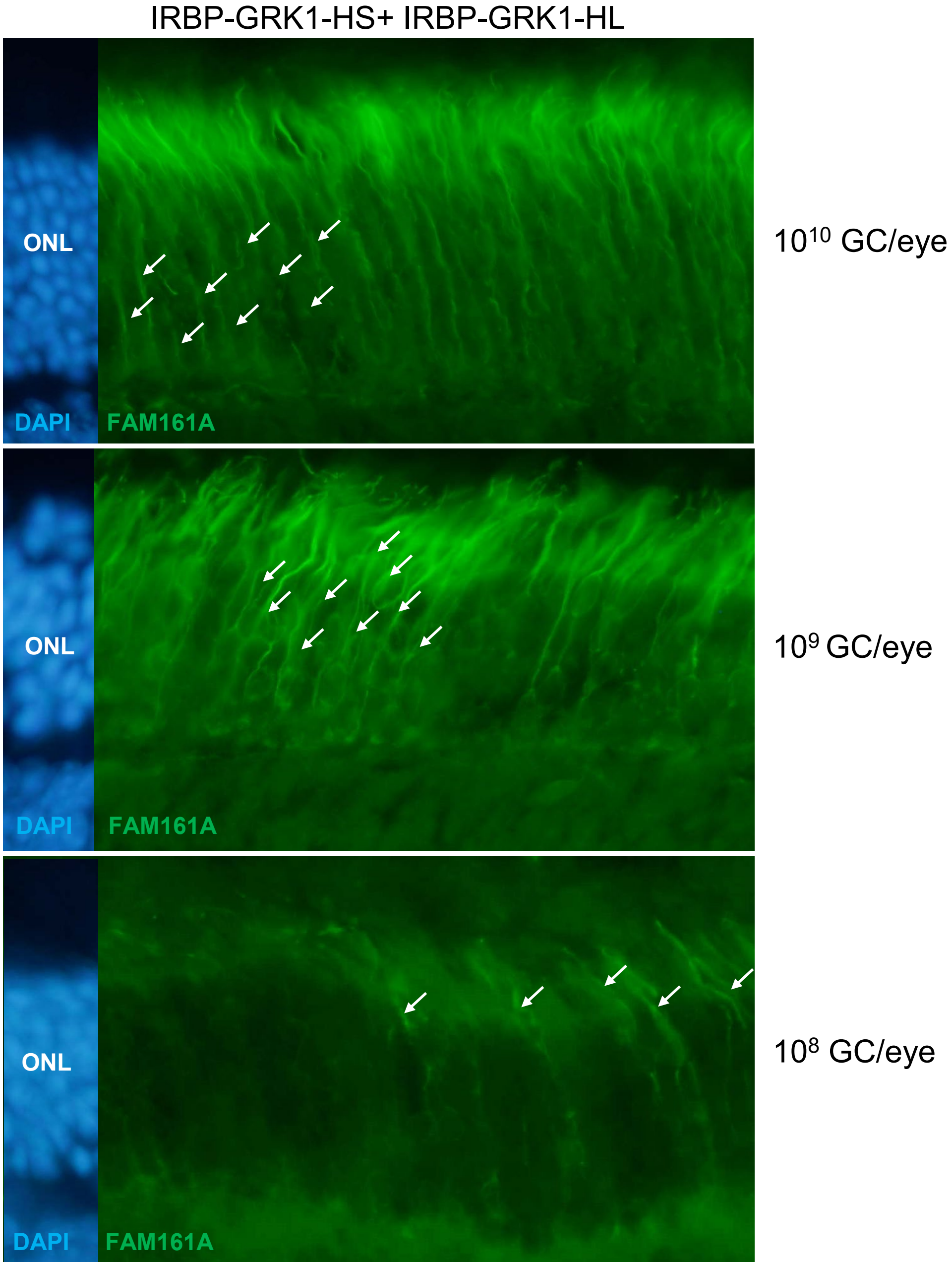


**Supplemental Figure 2. Spread expression in the photoreceptor driven by AAV2/8-IRBP-GRK1-FAM161A vectors is independent of the vector dose.** Subretinal injections of $10^8$, $10^9$, $10^{10}$ GC/eye of AAV2/8-IRBP-GRK1-FAM161A show the similar pattern of expression in the all photoreceptor body (arrows).

**Supplemental Figure 3**

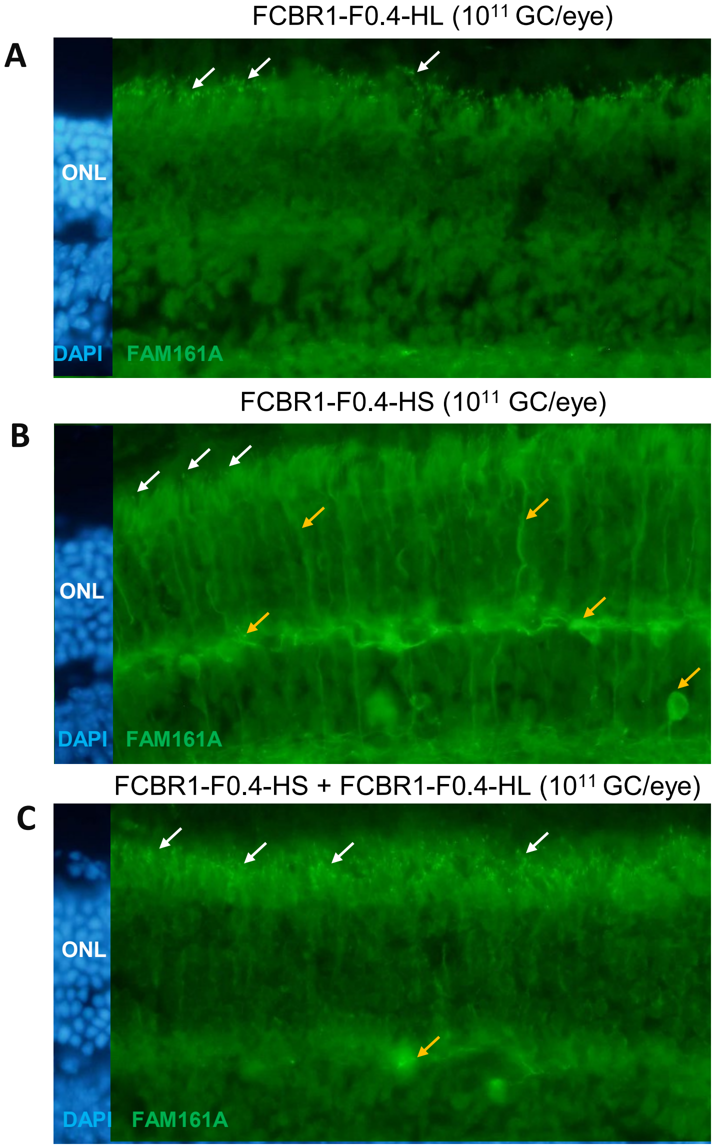


**Supplemental Figure 3. Expression pattern of FAM161A using AAV2/8-FCBR1-F0.4 vectors.** *Fam161a*$^{tm1b/tm1b}$ mice were subretinally injected with AAV2/8-FCBR1-F0.4-HL (**A**), AAV2/8-FCBR1-F0.4-HS (**B**) or AAV2/8-FCBR1-F0.4-HS + AAV2/8-FCBR1-F0.4-HL (**C**) vectors ($10^{11}$ GC/eye) at PN15 and FAM161A expression was analyzed 3 months later. Note for all conditions, the appearance of FAM161A-positive CC (white arrows), but the expression of the HS isoform induced ectopic expression of the protein (yellow arrows). (**C**) The co-injection of AAV2/8-FCBR1-F0.4-HS and AAV2/8-FCBR1-F0.4-HL produced a targeted and homogenous expression of the FAM161A in CC (white arrows) with also sometimes stronger labeling (yellow arrow). ONL: outer nuclear layer; HS: short isoform; HL: human long isoform; white arrows: correct labelling; orange arrows: modified labelling.

**Supplemental Figure 4**

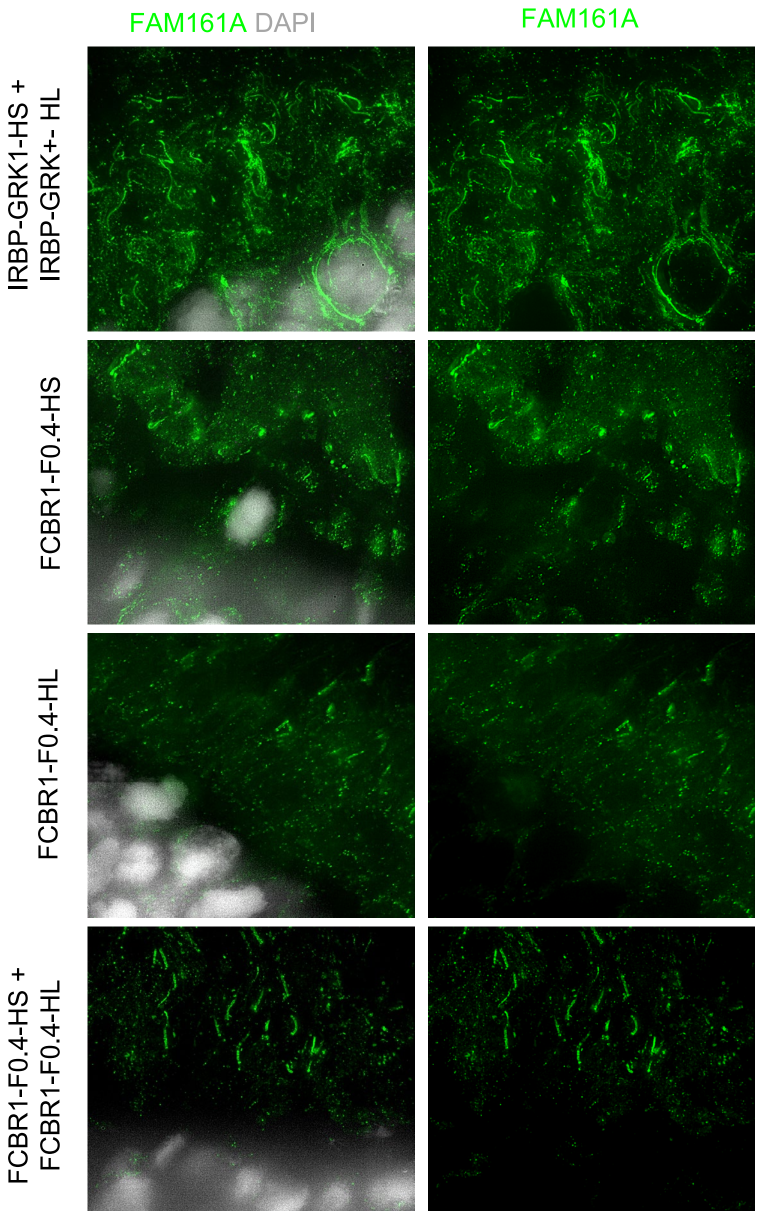


**Supplemental Figure 4. Abscence of ectopic expression pattern of FAM161A in the ONL after AAV2/8-FCBR1-F0.4-HS + AAV2/8-FCBR1-F0.4-HL treatment.** *Fam161a*$^{tm1b/tm1b}$ retina 3 month after treatment with AAV2/8-IRBP-GRK-HS + AAV2/8-IRBP-GRK-HL, AAV2/8-FCBR1-F0.4-HL, AAV2/8-FCBR1-F0.4-HS or AAV2/8-FCBR1-F0.4-HS + AAV2/8-FCBR1-F0.4-HL vectors ($10^{11}$ GC/eye) at PN15 are shown using Uex-M. Note the appearance of FAM161A labeling in the nucleus layer (white), with all treatments except with the combined injection with AAV2/8-FCBR1-F0.4-HS and AAV2/8-FCBR1-F0.4-HL.

**Supplemental Figure 5**

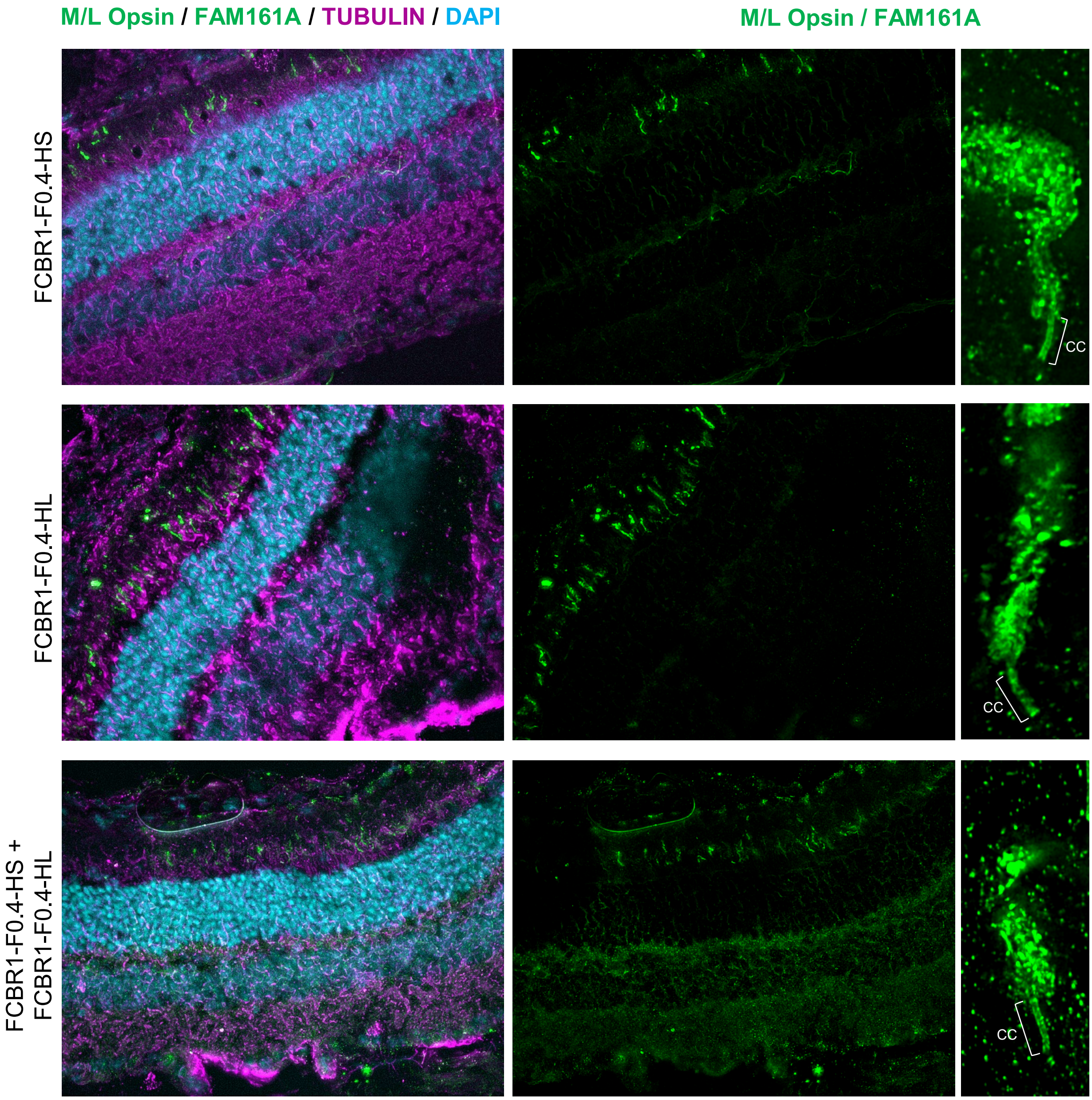


**Supplemental Figure 5. Restoration of FAM161A expression in the CC of cones.** *Fam161a*$^{tm1b/tm1b}$ retina 3 month after treatment with AAV2/8-FCBR1-F0.4-HS, AAV2/8-FCBR1-F0.4-HL or AAV2/8-FCBR1-F0.4-HS + AAV2/8-FCBR1-F0.4-HL vectors ($10^{11}$ GC/eye) at PN15 are shown using Uex-M. Note the appearance of FAM161A labeling in the cc of cones identified by labeling of their outersegment with M/L-opsin antibody (green).

**Supplemental Figure 6**

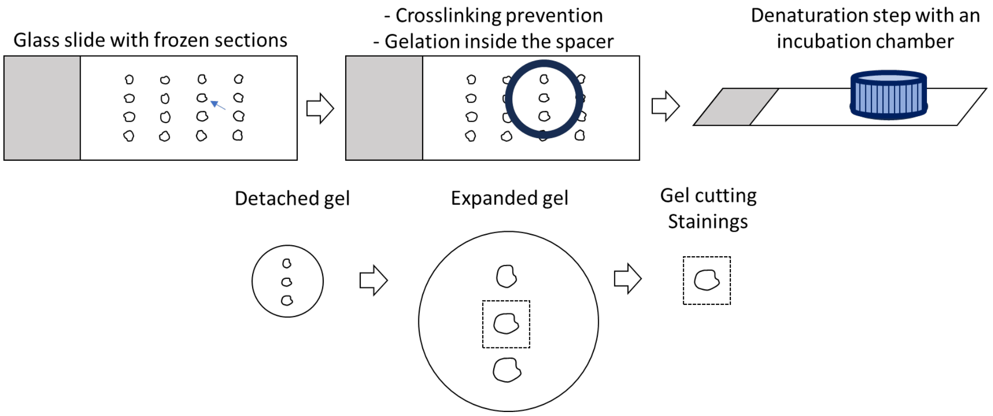


**Supplemental Figure 6. Flow chart of U-Ex-M process of retinal sections.**

**Supplemental Table 1. Primer sequences targeting the regulatory or coding regions included in the different plasmid constructed for the study.**

| | | |
|---|---|---|
| IRBP enhancer | forward | TGGAGGCAGAGGAGAAGG |
| IRBP enhancer | reverse | GCTTTATGAAGGCCAAAGAGG |
| hGRK1 promoter | forward | GGGCCCCAGAAGCCTGGTGG |
| hGRK1 promoter | reverse | GCCCTTGGCCTGTGGCCCG |
| FCBR1 | forward | GGGCTCAGACCTTAGAGACGGG |
| F0.4 | forward | TCCTCCATCAACTGTCTCCTTC |
| F0.4 | reverse | CGCCTCCGAGGCCTGAGC |
| FCBR2 | forward | CTCTCTCATTCTTTACAAATTCAGC |
| FCBR2 | reverse | TGATTTTGCACTGGATCCATTCTG |
| EFS | forward | GGCTCCGGTGCCCGTCAG |
| EFS | reverse | TCACGACACCTGTGTTCTGGCG |
| FAM161A cDNA | forward | ATGGCCACCTCCCACCG |
| FAM161A cDNA | reverse | TTGAAGAATCACACTGA |
| WPRE4 | forward | GGAATTCGAGCATCTTACCG |
| WPRE4 | reverse | CTTCCCCGACAACACCAC |

**Supplemental Table 2. Antibodies used in the study.**

| Antigen | Use | Antibody name | Reference |
|---|---|---|---|
| FAM161A | WB, ICC, IHC | HPA032119 | Sigma |
| FAM161A | Uex-M | PA556935 | Thermo Fischer Scientific |
| GADPH | WB | MAB74 | Chemicon |
| GFP | ICC | Ab290 | Abcam |
| GFP | IHC | Ab1218 | Abcam |
| Acetylated-TUBULIN | ICC | T7451 | Sigma |
| $\alpha$-TUBULIN | Uex-M | AA345 - scFv-F2C | ABCD antibodies |
| $\beta$-TUBULIN | Uex-M | AA344 - scFv-S11B | ABCD antibodies |
| Rhodopsin | IHC, Uex-M | MA5-11741 | Thermo Fischer Scientific |
| M/L-opsin | Uex-M | AB 5405 | Millipore |
| POC5 | Uex-M | A303-341A | Bethyl |
| CEP290 | Uex-M | 22490-1-AP | Proteintech |
| LCA5 | Uex-M | 19333-1-AP | Proteintech |
| IFT81 | Uex-M | 11744-1-AP | Proteintech |
| Anti-rabbit Alexa Fluor 488 | ICC, IHC | A11070 | Thermo Fischer |
| Anti-mouse Alexa Fluor 633 | ICC, IHC | A21053 | Thermo Fischer |
| anti-Rabbit IgG HRP | WB | NA0934 | Amersham Biosciences |
| anti-Mouse IgG HRP | WB | NA0931 | Amersham Biosciences |
